# Altered PBP4 and GdpP functions synergistically mediate MRSA-like high-level, broad-spectrum β-lactam resistance in *Staphylococcus aureus*

**DOI:** 10.1101/2023.10.26.564222

**Authors:** Li-Yin Lai, Nidhi Satishkumar, Sasha Cardozo, Vijay Hemmadi, Leonor B. Marques, Liusheng Huang, Sergio R. Filipe, Mariana G. Pinho, Henry F. Chambers, Som S. Chatterjee

## Abstract

Infections caused by *Staphylococcus aureus* are a leading cause of mortality worldwide. *S. aureus* infections caused by Methicillin-Resistant *Staphylococcus aureus* (MRSA) are particularly difficult to treat due to their resistance to Next Generation β-lactams (NGB) such as Methicillin, Nafcillin, Oxacillin etc. Resistance to NGBs, which is alternatively known as broad-spectrum β- lactam resistance is classically mediated by PBP2a, a Penicillin-Binding Protein encoded by *mecA* (or *mecC*) in MRSA. Thus, presence of *mec* genes among *S. aureus* serves as the predictor of resistance to NGBs and facilitates determination of the proper therapeutic strategy for a staphylococcal infection. Although far less appreciated, *mecA* deficient *S. aureus* strains can also exhibit NGB resistance. These strains, which are collectively termed as Methicillin-Resistant Lacking *mec* (MRLM) are currently being identified in increasing numbers among natural resistant isolates of *S. aureus*. The mechanism/s through which MRLMs produce resistance to NGBs remains unknown. In this study, we demonstrate that mutations that alter PBP4 and GdpP functions, which are often present among MRLMs can synergistically mediate resistance to NGBs. Furthermore, our results unravel that this novel mechanism potentially enables MRLMs to produce resistance towards NGBs at levels comparable to that of MRSAs. Our study, provides a fresh new perspective about alternative mechanisms of NGBs resistance, challenging our current overall understanding of high-level, broad-spectrum β-lactam resistance in *S. aureus*. It thus suggests reconsideration of the current approach towards diagnosis and treatment of β-lactam resistant *S. aureus* infections.

## Introduction

*Staphylococcus aureus* is a Gram-positive bacterium that is a frequent colonizer of the human population. It is also an opportunistic pathogen with the potential to cause a range of infections including skin and soft tissue infections, bacteremia, osteomyelitis and sepsis (1). Along with its pathogenicity, *S. aureus* is infamous for its ability to evade antibiotic treatment either in the form of drug resistance or tolerance (2, 3). The ability of *S. aureus* to cause pathogenic infections, coupled with its potential to evade the actions of antibiotics, makes it one of the leading causes of morbidity in the world (4). Due to their high safety and efficacy, β-lactams are one of the most successful and commonly prescribed classes of antibiotics for bacterial infections (5). β-lactams target the bacterial cell wall by binding and inactivating a class of proteins involved in the cell wall synthesis and maintenance, known as Penicillin-Binding Proteins (PBPs) that result in the weakening of the bacterial cell wall and subsequent cell death (6). Resistance to β-lactams can classically occur either as narrow-spectrum resistance associated with BlaZ, a β-lactamase that hydrolyzes and inactivates the β-lactam ring of the drug (7), or as broad-spectrum resistance attributed to *mecA* (or *mecC*), a gene that encodes PBP2a, which has decreased affinity to β- lactams (8). While narrow-spectrum resistance is limited to early generation drugs such as Penicillin, broad-spectrum resistance renders the entire class of β-lactam drugs ineffective, including Next Generation β-lactams (NGBs). This reduced affinity towards β-lactams enables bacteria to survive in presence of antibiotics, providing high-level resistance in strains characterized as Methicillin Resistant *S. aureus* (MRSA) (9), which resulted in over 100,000 deaths in 2019 in the US alone (10). The subsequent development of Ceftaroline, an advanced-generation NGB with a high affinity for PBP2a has largely aided in combating *mecA*-associated β-lactam resistance (11).

NGB resistance can also be detected in *S. aureus* isolates that do not contain *mec* genes, referred to as Methicillin-Resistant Lacking *mec* (MRLM) strains. Although first detected in the 1980s, the underlying mechanism/s of resistance in MRLMs remain unknown (12). A recent spur in detection of natural MRLM isolates prompted efforts to map the basis of NGBs resistance among MRLMs (13-15). These studies, which primarily employed whole genome sequencing of MRLM isolates, identified mutations associated with *pbp*s or *gdpP* in high frequencies.

Prior to these recent efforts, our group performed a series of laboratory passaging experiments with NGB sensitive strains of *S. aureus* that lacked *mecA* with the aim to determine non-classical mechanisms of β-lactam resistance that are independent of PBP2a or BlaZ (16-18). Passaging of strains in NGBs produced variants that displayed high-level, MRSA-like resistance. These resistant strains most prominently harbored mutations in *pbp4* (encodes for PBP4) and *gdpP* (encodes for <u>G</u>GDEF <u>d</u>omain <u>p</u>rotein containing <u>p</u>hosphodiesterase), suggesting that the mutations associated with these genes played an important role in facilitating high-level NGB resistance (19). Our subsequent findings demonstrated that *pbp4*, through regulatory site and/or gene associated mutations resulted in β-lactam resistance (19, 20). On the other hand, mutations associated with the catalytic domain of *gdpP*, a phosphodiesterase that cleaves the second messenger cyclic-di-AMP (CDA), led to increased amounts of CDA within the cell, suggesting that the mutations led to GdpP’s loss-of-function (21). Further, the *gdpP* mutants evaded β-lactam treatment by drug tolerance, a phenomenon where bacteria can survive β-lactam challenges without resulting in a change in MIC (21, 22). However, altered functioning of neither PBP4, nor GdpP, on their own accounted for the high-level, broad-spectrum resistance to β-lactams that we reported previously among the laboratory-passaged NGB resistant strains (16, 18).

In this study, we examined the combined effect of mutations associated with *pbp4* and *gdpP* on resistance by employing MIC assay, population analysis and growth assay, along with determining their effect on tolerance by performing a Tolerance-Disk test (TD test) (23). Strains containing mutations associated with both *pbp4* and *gdpP* had significantly increased survival when challenged with different NGBs such as Nafcillin, Oxacillin and Ceftaroline, demonstrating high-level, broad-spectrum β-lactam resistance. Alterations in only PBP4 or GdpP did not result in the high-level resistant phenotype, suggesting a synergistic effect between the two mechanisms. This synergistic action on resistance provides an explanation for the high-level resistance seen in the aforementioned laboratory-passaged strains. These findings thus suggest that MRLM strains, by means of altering PBP4 and GdpP functions, can develop resistance to NGBs to the extent that is comparable to MRSA strains. *In vivo* studies performed with *C. elegans* demonstrated that alterations in PBP4 and GdpP allows for successful infection in presence of β- lactams, thus also having the potential to lead to therapy failure.

## Materials and Methods

### Bacterial strains

*S. aureus* strains were cultured in Tryptic soy broth (BD Biosciences, USA) media with aeration or on Tryptic soy agar (BD Biosciences, USA) plates at 37°C. Strains carrying the *pTX*_Δ_ plasmids were grown in media containing 12.5 mg/L tetracycline. The strains used in this study are listed in **Table S1**.

### Construction of mutants

Primers used in this study are listed in **Table S2**. Mutations in the SF8300ex strain were introduced through allelic replacement as previously described (24). Briefly, the 1 kb up- and downstream regions of the target gene (*gdpP* or *pbp4*) were amplified and splice-overlap extension PCR (SOE PCR) was performed. The PCR product was cloned into either the *pKOR1* plasmid (25) or *pJB38* plasmid (26) and the resultant plasmid was confirmed by sequencing (Eurofins Genomics, USA) before its transformation into RN4220, following which it was introduced into the recipient strain. Standard allelic replacement procedure was then carried out as previously described (25) and the obtained mutant was validated using PCR and/or Sanger sequencing.

### Minimum Inhibitory Concentration (MIC) Assay

MICs were determined by broth microdilution method as described previously (27). Briefly, 1x10^5^ CFU bacteria were incubated for 48h at 37°C in 0.2 mL cation-adjusted Mueller-Hinton broth (CAMHB) (BD Biosciences) containing increasing concentrations of antibiotic (0.25 mg/L to 256 mg/L for Nafcillin and Oxacillin, 0.25 mg/L to 4 mg/L for Ceftaroline). MIC was recorded as the lowest concentration without growth at 48h. MIC assay was performed twice in order to ensure reproducibility.

### Population assay

Population assay was performed as previously described (27). Briefly, *S. aureus* strains were cultured in 3 mL TSB at 37°C overnight. A 10 µL volume of serially diluted bacterial culture was spotted onto the prepared antibiotic-containing TSA plates and incubated at 37°C for 48h. Plates were read and expressed as CFU mL^−1^. Tetracycline (12.5 mg/L) was added to media and plates for the complemented strains carrying the *pTX*_Δ_ plasmid. Population assay was performed twice to ensure reproducibility.

### Growth curve assay

Growth assays were performed as previously described using Bioscreen C, an automated growth curve analysis system (Growth Curves USA) (28). Briefly, overnight cultures of bacteria were diluted to an OD_600nm_ of 0.1 in TSB with or without antibiotics and 200 µL of the bacterial dilution was pipetted into each well of a Bioscreen C plate in triplicates. Growth curves were performed with continuous orbital shaking for a period of 12h at 37°C. The data was then analyzed for OD_600nm_ values at each time point for every antibiotic concentration used. Growth assay was performed twice in triplicates to ensure reproducibility, except for Ceftraroline, it being expensive and hard to acquire.

### Tolerance Disk Test (TD-test)

TD-test was carried out as previously described with minor modifications (29). Briefly, overnight bacterial cultures were adjusted to OD_600nm_ of 0.1, and 100 µL bacteria were plated onto TSA plates. A 6-mm disks containing 1 µg Nafcillin (or Oxacillin) was placed in the bacterial lawn, and the plates were left to incubate at 37°C. After 18h, the zone of inhibition for each plate was marked and the antibiotic-containing disks were replaced with 4 mg glucose-containing disks. The plates were incubated at 37°C for another 2 days, following which tolerant colonies were enumerated. Colonies were counted as described in **Fig S5C**, where the inner half of the zone of inhibition was annotated, and the colonies present within this region were counted as tolerant. TD-test was performed twice to ensure reproducibility.

### Immunoblotting

Overnight cultures of bacteria were subcultured in 50 mL flasks containing TSB such that the initial OD_600nm_ of the flasks was 0.1. The cells were cultured to OD_600nm_ = 1, following which they were collected and resuspended in PBS containing CompleteMini protease inhibitor cocktail (Roche). The cells were mechanically lysed using the FastPrep (MP Biochemicals) and whole cell lysates were obtained. The cell membrane fraction was isolated from the lysates by performing ultracentrifugation at 66000g for 1h (Sorvall WX Ultra 80 Centrifuge, Thermo Fisher Scientific). After resuspending the obtained pellet with PBS, protein estimation was carried out using the Pierce BCA Protein Assay kit (Thermo Fisher). The samples were separated by performing SDS-PAGE on a 10% gel, following which they were transferred onto a low-fluorescence PVDF membrane (Millipore). Blocking was performed for 1h (5% skimmed milk in Tris-buffered saline containing 0.5% Tween) and primary antibody staining was carried out overnight at 4°C (polyclonal anti-PBP2a, custom antibody from Thermo Fisher, 1:1000 or polyclonal anti-Sortase A, custom antibody from Thermo Fisher 1:1000). Secondary antibody staining was performed using an anti-rabbit antibody (Azure anti-rabbit NIR700 or Azure anti-rabbit NIR800 at 1:20000 dilution). The blots were imaged using the Azure C600 imager.

### β-lactamase assay

Nitrocefin discs (BD BBL™ Cefinase™ β-Lactamase Detection Discs) were placed onto tryptic soy agar (TSA) plates following which bacteria were streaked onto the discs using sterile toothpicks. The plates were incubated at 37°C for 30 min, following which, the plates were observed for change of color. A color change to red was considered as β-lactamase positive.

### Intracellular CDA measurement from bacteria

CDA measurement was carried out as previously described with a few modifications (21, 30). Briefly, bacterial cells were collected after 6h of culture, washed, and lysed in phosphate-buffered saline (1X PBS) containing 1 mM EDTA using a FastPrep-24 homogenizer (MP Biomedicals). Bacterial cytosolic fraction was collected upon centrifugation. Aliquots of 40 μL cytosolic samples were mixed with 10 μL internal standard (20 ng/mL tenofovir, a phosphate group containing compound that is negatively charged in solution, similar to CDA), and 10 μL of the resulting mixture was injected into a liquid chromatography-tandem mass spectrometer (LC-MS/MS) system. The LC-MS/MS system consisted of AB Sciex API 5000 tandem mass spectrometer, Shimadzu Prominence 20ADXR ultra-fast liquid chromatography (UFLC) pumps, and SIL- 20ACXR autosampler. A Hypercarb analytical column (100 by 2.1 mm, 3 mm; Thermo Fisher Scientific, USA) and mobile phases 100 mM ammonium acetate (Buffer A, pH 9.8) and acetonitrile (Buffer B) were used for separation. Electrospray ionization in negative ion mode as the ion source and multiple reactions monitoring with ion pairs m/z 657/133 for CDA and m/z 286/133 for the internal standard were used for quantification. Calibration standards were prepared with synthetic CDA (Biolog) dissolved in the PBS containing 1 mM EDTA. The calibration range was 10 to 500 ng/mL with lower limit of quantification at 10 ng/mL.

### Bocillin assay

Bocillin assay was performed as described in our previous study with modifications. Briefly, *S. aureus* strains were cultured in TSB, and cell density was adjusted to OD_600nm_ = 0.1 in 50 mL cultures. Samples were collected when the bacterial cells reached an OD_600nm_ of 1. The lysates were labeled with 1 μM Bocillin-FL (Thermo Fisher Scientific) at 35°C for 30 min, and the reaction was terminated by adding a sample buffer and boiling the samples for 10 mins. 20 μg of each sample was analyzed by 10% SDS-PAGE. The gel was scanned with Typhoon 9410 imager (Amersham/GE Healthcare) to visualize the Penicillin Binding Proteins on the gel.

### Bacterial peptidoglycan purification and analysis

Muropeptide purification from *S. aureus* was performed as previously described (31, 32). Briefly, cells were harvested at the exponential phase (OD_600nm_ 0.8-0.9) and boiled in 4% SDS to inactivate cell wall modifying enzymes, following which the SDS was removed by washing cells several times with water. Cells were then lysed mechanically using SpeedMill PLUS homogenizer, following which cell wall, and subsequently peptidoglycan purification was performed as described previously (31, 32). Purified peptidoglycan was treated with Mutanolysin to obtain muropeptides that were reduced with sodium borohydrate in borate buffer and analysed by reverse phase HPLC using a Waters Acquity CSH C18 column. Elution was performed using a gradient up to 80% acetonitrile (0.3 mL/ min) for 28 minutes at 205nm.

### Caenorhabditis elegans killing assay

Infection of *C. elegans* DH26 was performed as described previously (28). Briefly, after age synchronization, 15 L4-young adult worms were incubated with 1.5 x 10^6^ *S. aureus* in each well of a 96-well flat plate. Bacteria were added in 100 µL liquid assay media (80% M9 buffer, 20% TSB, 10 mg/L cholesterol, and 7.5 mg/L nalidixic acid) following which Nafcillin was added at the indicated concentrations. Infection studies were conducted at 26°C for 72h, and the survival rate was recorded.

*C. elegans* gut CFU determination was performed as previously described (28, 33). Briefly, 100 µL of M9 buffer containing 10 mM sodium azide and 100 mg/L gentamicin (hereinafter buffer 1) was transferred to 96-well plates. Worms were transferred to 1.5 mL micro-centrifuge tubes and washed twice with 500 µL buffer 1. After gentamicin treatment, worms were washed three times with M9 buffer containing 10 mM sodium azide (hereinafter buffer 2) to remove gentamicin. Worms were then suspended in 300 µL buffer 2. 50 µL supernatant (without worms) was removed, and the remaining 250 µL sample (with worms) was lysed by vortexing with 200 mg of zirconium beads for 2-5 min. The before and after lysis samples were plated on TSA plates and incubated at 37°C overnight. CFU determination was carried out the following day.

### Hemolysis assay

Overnight cultures of bacterial strains were spotted onto TSA-blood plates (5% sheep blood) in increasing volumes (2 µL, 5 µL, 10 µL) as indicated, and were incubated at 37°C overnight, following which the plate was stored at 4°C before recording the hemolysis pattern.

### Statistical Analysis

Statistical significance was analyzed using a one-way analysis of variance (ANOVA) or by a two-tailed Student’s *t*-test on the GraphPad Prism software (version 8, La Jolla, CA, USA).

## Results

### Mutations associated with *pbp4* and *gdpP* are detected in high frequencies among MRLM strains

Our previous studies targeted at identifying non-classical mechanisms of NGB resistance in *S. aureus* demonstrated that mutations associated with *pbp4* and *gdpP* occurred in high frequencies in strains lacking *mecA* when passaged in NGBs (16-18). Among the six NGB-passaged strains studied, *pbp4*-associated mutations were detected in five strains whereas *gdpP*-associated mutations were detected in all strains **(Fig S1, Supplemental file 1)**. The *pbp4*-associated mutations were either regulatory site associated, i.e. mutations located upstream of the *pbp4* start codon, and/or were missense mutations detected within the gene. *pbp4* regulatory site-associated mutations led to increased expression of PBP4, resulting in a highly cross-linked cell wall (24) while the missense mutations either led to altered protein structure, which lowered drug affinity or led to increased thermal stability that resulted in resistance (34, 35). Mutations associated with *gdpP* led to partial or complete loss of its phosphodiesterase activity (referred to as loss-of-function mutations) causing an increase in CDA concentrations in bacteria that enabled cells to survive in high concentrations of β-lactams- a phenomenon commonly referred to as antibiotic tolerance (36).

Following the detection of mutations in laboratory-passaged strains, we performed a literature search to determine if mutations associated with *pbp4* and/or *gdpP* were also relevant in naturally isolated MRLM strains. Indeed, three independent studies aimed at characterizing mutations associated with MRLM isolates each reported the presence of similar mutations associated with *pbp4* and/or *gdpP* (13-15) **(Fig 1, Fig S2, Supplemental file 1)**. The criteria used by all three studies to classify an isolate as MRLM, namely, the absence of *mec*-genes, and phenotypic resistance to Oxacillin (MIC > 2mg/L) and/or Cefoxitin (MIC > 4mg/L) were consistent. This allowed us to uniformly classify the isolates based on their mutations, predict their phenotypic manifestations and their subsequent effect on β-lactam resistance despite being reported in independent studies.

**Figure 1.**
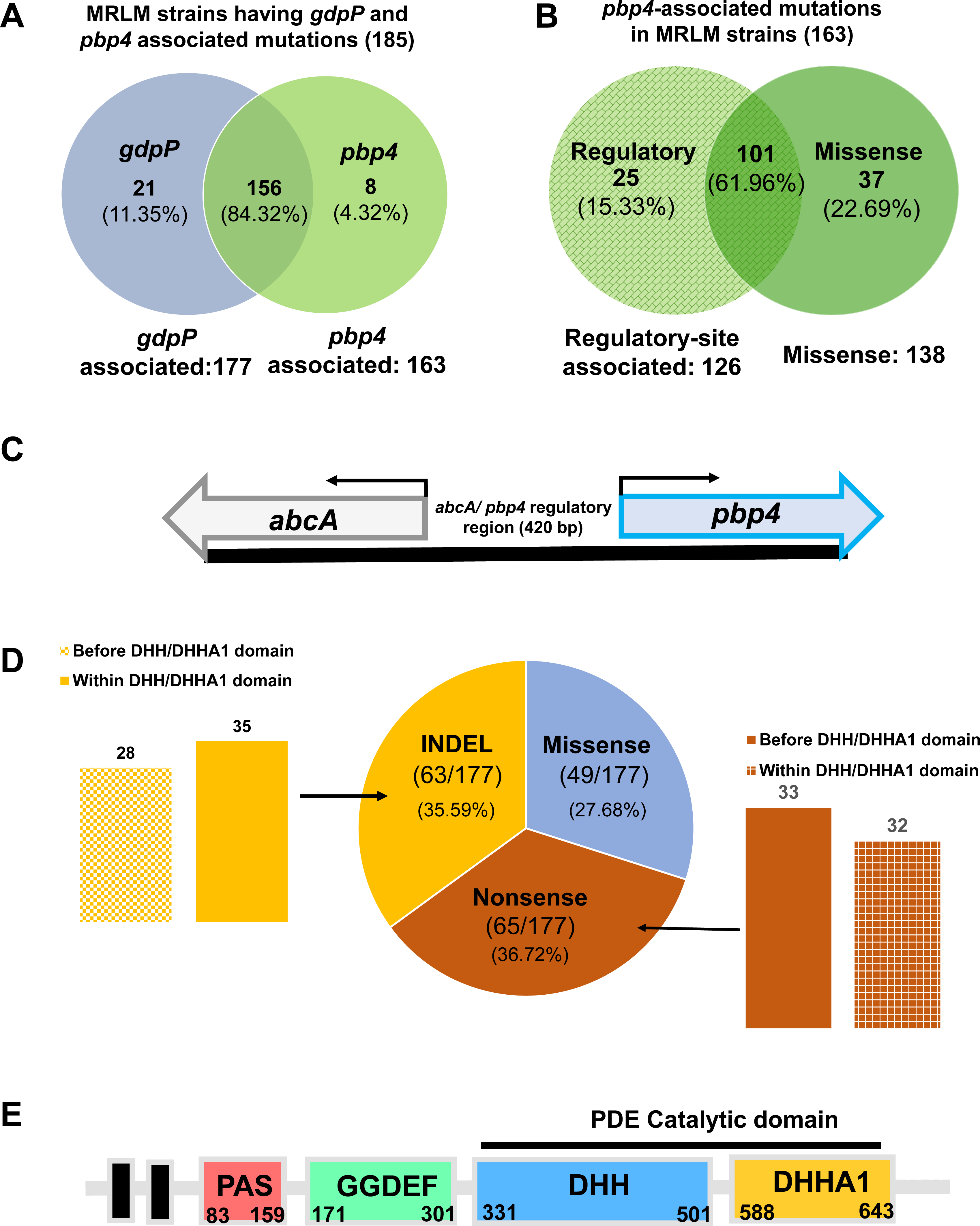
*GdpP* and PBP4 mutations that were identified in MRLM isolates. **(A)** Venn diagram describing occurrence *pbp4* and/or *gdpP* mutations in MRLM strains described in previous studies. **(B)** Venn diagram describing the occurrence of *pbp4* regulatory site associated and/or missense mutations in MRLM strains described previously **(C)** Schematic diagram of *pbp4* and *abcA* genes separated by a 420 bp intergenic, regulatory site region. **(D)** Illustration and distribution of *gdpP*-associated mutations (INDELs, missense and nonsense) detected in MRLM isolates. *gdpP*-associated mutations were further classified based on location of mutation with respect to the DHH/DHHA1 catalytic domain. **(E)** Schematic representation of different domains in the *gdpP* gene. The phosphodiesterase catalytic activity of GdpP is mediated by its DHH/DHHA1 domains.

Of the isolates reported in these studies, a total of 156 MRLM isolates (84.32%) had mutations associated with both *pbp4* and *gdpP*, while a relatively smaller subset of isolates had mutations associated with only *pbp4* (8 isolates, 4.32%) or only *gdpP* (21 isolates, 11.35%) **(Fig 1A)**. *pbp4*- associated mutations were either regulatory site-associated (25 of 163, 15.33%), missense mutations, that resulted in amino acid substitutions in the *pbp4* gene (37 of 163, 22.69%), or detected as both, regulatory site and missense mutations (101 of 163, 61.96%) **(Fig 1B**, **Fig 1C)**. A majority of *pbp4*-associated mutations were thus detected in both, the regulatory site as well as the gene. 65 of the 177 (36.72%) isolates that possessed *gdpP* mutations were nonsense mutations, causing premature truncation of GdpP due to the introduction of a stop codon **(Fig 1D)**. 63 isolates (35.59%) were INDELs resulting in frameshift mutations, while 49 isolates (27.68%) had missense mutations. Importantly, our analysis led to the observation that a large proportion of the mutations (128 out of 177) would result in loss of GdpP function due to premature truncation either upstream of its catalytic DHH/DHHA1 domains (33 isolates with non-sense mutations, 28 with INDELs) or within the DHH/DHHA1 domains (32 isolates with non-sense mutations, 35 with INDELs) **(Fig 1D**, **Fig 1E)**. This indicated that a large portion of the isolates would display increased CDA concentration, as well as β-lactams tolerance, a phenotype that was observed previously among our resistant laboratory-passaged strains (21).

### Mutation-associated altered functions of PBP4 and GdpP synergistically produced high-level, broad-spectrum NGB resistance

Of the aforementioned laboratory-passaged strains, CRB was the strain where this unusual mode of resistance was initially identified (17). CRB contained mutations associated with *pbp4* (regulatory site and missense), and *gdpP* (loss-of-function). (16, 17). Unlike the other passaged strains, CRB did not have mutations associated with any other *pbps*, and displayed high-level β- lactam resistance with a MIC of 256 mg/L for Nafcillin (compared to a MIC of 1 mg/L for its susceptible parent), thus underscoring the potential roles of PBP4 and GdpP in high-level NGB resistance (16, 18). Further probing of the effect of *pbp4*-associated mutations was performed by introducing the regulatory site mutations (a 36 bp duplication 290 bp upstream the *pbp4* start codon) and missense mutations (E^183^A and F^241^R) detected in CRB into a NGB-sensitive parental strain. The presence of *pbp4*-associated mutations led to a considerable increase in resistance as seen by a MIC of 4 mg/L for Nafcillin, when compared to its NGB-sensitive parental strain (19). While this increase in resistance was significant, it was not as high as that displayed by CRB, where the increase in MIC was 256-fold (16). CRB also contained loss-of-function mutations in *gdpP*. Loss-of-function mutations in *gdpP* did not result in a change in MIC, but instead led to NGB-tolerance (21). Taken together, *pbp4* and *gdpP* mutations resulted in broad-spectrum β- lactam resistance and tolerance respectively; however, neither of them could independently produce the high-level resistant phenotype that was detected in CRB (16, 19). This led us to hypothesize that functional alteration of both PBP4 and GdpP are together required for high-level broad-spectrum β-lactam resistance. We thus sought to assess the effects of alterations in both, PBP4 and GdpP by studying the effect of mutations detected in CRB (16, 17). The *pbp4* regulatory site (P*pbp4*\*) and missense (*pbp4*\*\*) mutations were introduced in SF8300ex (Wtex); a *mecA* and *blaZ* excised wild-type (Wt) SF8300, a USA300, a prominent community-associated strain **(Table S1, Fig 2A)** (2, 19, 28). Additionally, the loss-of-function effect of *gdpP* mutations was introduced by deletion of *gdpP*, thus resulting in the strain Wtex P*pbp4*\* *pbp4*\*\* Δ*gdpP*, or, the triple mutant. The resultant isogenic strains allowed us to assess the role of PBP4 and *gdpP* mutations together **(Fig 2A)** in comparison to their parental strain (Wtex) as well as the wild-type MRSA strain (Wt). At first, immunoblotting for PBP2a (76.1 kDa), the gene product of *mecA* was carried out to determine the accurate phenotypes of the obtained isogenic mutants. As expected, PBP2a was only detected in Wt **(Fig 2B)** indicating that it was absent in all Wtex background strains. Further, a nitrocefin disc assay was performed to determine β-lactamase activity of the strains, which also confirmed that only the Wt strain contained BlaZ **(Fig 2C)**. The above tests thus verified that the studied mutants were devoid of the classical mediators of β-lactam resistance. Phenotypes associated with *pbp4* mutations were verified through Bocillin assay **(Fig 2D, Fig S3)**. Strains with regulatory site mutations (P*pbp4*\*), had an increased expression of PBP4 compared to all other strains. Alanine substitution of the essential active site serine of PBP4, S^75^A, did not allow for Bocillin binding and thus no PBP4 bands were detected in the corresponding strains **(Fig 2D)**. Deletion of *gdpP* was verified by measuring levels of intracellular CDA; elevated levels of CDA was detected in strains that lacked *gdpP* compared to the other isogenic strains, confirming the Δ*gdpP* phenotype in these mutants **(Fig 2E).**

**Figure 2.**
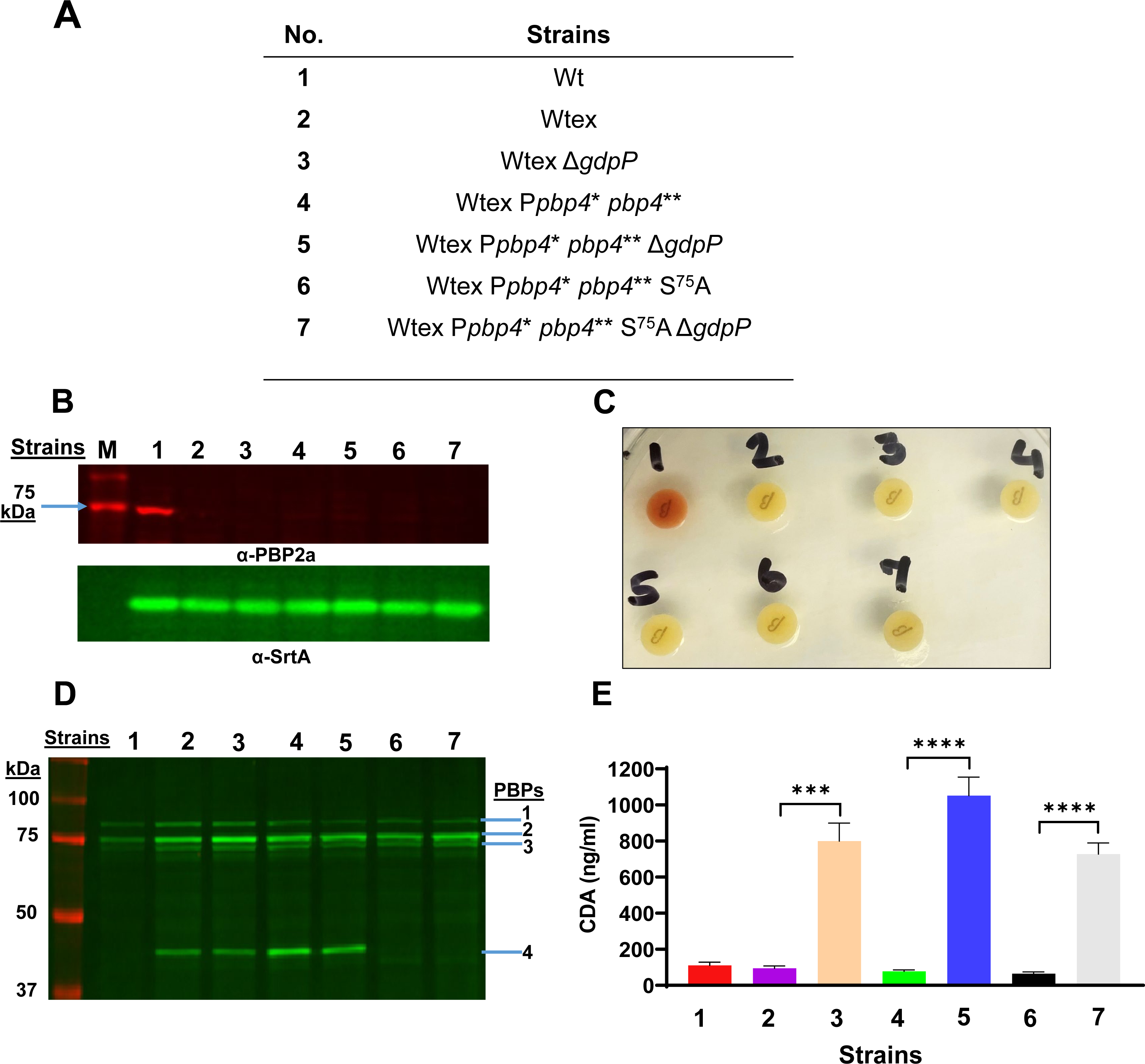
Phenotypic characterization of the strains used in this study. **(A)** List of strains used for obtaining the results for sub figures B-E. **(B)** Immunoblotting to detect PBP2a (α-PBP2a, 76.1 kDa, upper panel) was present in only lane 1 (Wt) and Sortase-A (α-SrtA, lower panel) as a loading control that was detected for all samples. “M” represents Protein Molecular Weight Marker. **(C)** Nitrocefin disk test indicated that only Wt had β-lactamase activity, as seen by the color change of the disk from yellow to red. Color of the disks streaked with other strains remained unchanged. **(D)** Bocillin assay to detect PBPs 1 through 4. Strains with *pbp4* regulatory site mutations (lanes 4 and 5) showed enhanced PBP4 expression compared to the isogenic Wtex strain (lane 2). Strains with the *pbp4* S^75^A mutation (lanes 6 and 7) did not show any PBP4 band. Δ*gdpP* (lanes 5 and 7) did not have an effect on expression of PBPs. PBP expression levels were visualized by the Typhoon 9410 imager (Amersham/GE Healthcare). **(E)** Measurement of intracellular levels of CDA for the studied strains. Strains with the Δ*gdpP* mutation show high levels of CDA compared to their isogenic pairs. P-value = 0.0003 for Wtex versus Wtex Δ*gdpP*; P-value < 0.0001 for Wtex P*pbp4*\* *pbp4*\*\* versus P*pbp4*\* *pbp4*\*\* Δ*gdpP*, and Wtex P*pbp4*\* *pbp4*\*\* S^75^A versus Wtex P*pbp4*\* *pbp4*\*\* S^75^A Δ*gdpP*.

Following the validation of strain phenotypes, we assessed their resistance profiles. MIC assay for P*pbp4*\* *pbp4*\*\* Δ*gdpP*, the triple mutant, revealed a drastic increase in resistance to Nafcillin and Oxacillin (64 fold), along with Ceftaroline (8 fold) when compared to Wtex **(Table 1)**. This increase in MIC values seen in the triple mutant was significantly higher than Wtex Δ*gdpP* as well as Wtex P*pbp4*\* *pbp4*\*\*, suggesting a synergistic role of altered functions of PBP4 and GdpP **(Table 1)**. Validation of this synergism was performed either by including the *pbp4* non-functional mutation, S^75^A in the triple mutant (Wtex P*pbp4*\* *pbp4*\*\* S^75^A Δ*gdpP)*, or, by complementing it with a functional GdpP (Wtex P*pbp4*\* *pbp4*\*\* Δ*gdpP* [*gdpP*]), both of which resulted in susceptibility to the selected NGBs **(Table 1 and Table 2)**. Additionally, the increase in MIC for the triple mutant was comparable to the *mecA*-containing Wt strain, and in case of Ceftaroline, the MIC was 4-fold higher than that of Wt. These findings thus demonstrated that alterations in PBP4 and GdpP could synergistically result in MRSA-like, high-level NGB resistance. Population assay validated these findings, as the triple mutant had significantly increased resistance to Nafcillin and Oxacillin compared to the parent strain, Wtex, and could survive in high concentrations of β-lactams similar to Wt **(Fig 3A, 3B)**. In alignment with the MIC assay values, the triple mutant had increased survival compared to Wtex Δ*gdpP* and Wtex P*pbp4*\* *pbp4\*\**, reiterating the synergistic actions of altered PBP4 and GdpP. The inclusion of the S^75^A mutation in the triple mutant (Wtex P*pbp4*\* *pbp4*\*\* S^75^A Δ*gdpP*) resulted in absolute susceptibility to Nafcillin and Oxacillin **(Fig 3)**. Similar resistant phenotypes were observed when a growth assay was performed; in absence of NGBs, strains with Δ*gdpP* had a growth defect, a well-established phenotype associated with the mutation. However, Wt and Wtex P*pbp4*\* *pbp4*\*\* Δ*gdpP* were the only strains that survived in presence of Nafcillin and Oxacillin **(Fig S4)**. Complementation with functional GdpP (Wtex P*pbp4*\* *pbp4*\*\* Δ*gdpP* [GdpP]) also led to susceptibility in a population analysis **(Fig 4A and 4B).** GdpP-complementation was phenotypically verified by measuring intracellular CDA levels, where the mutant complemented with a functional GdpP contained decreased amounts of CDA compared to the mutants complemented with an empty vector, [E] **(Fig 4C)**. The high-level, MRSA-like resistance to various NGBs was thus attributed to a synergistic effect brought about by mutations in both *pbp4* and *gdpP*.

**Figure 3:**
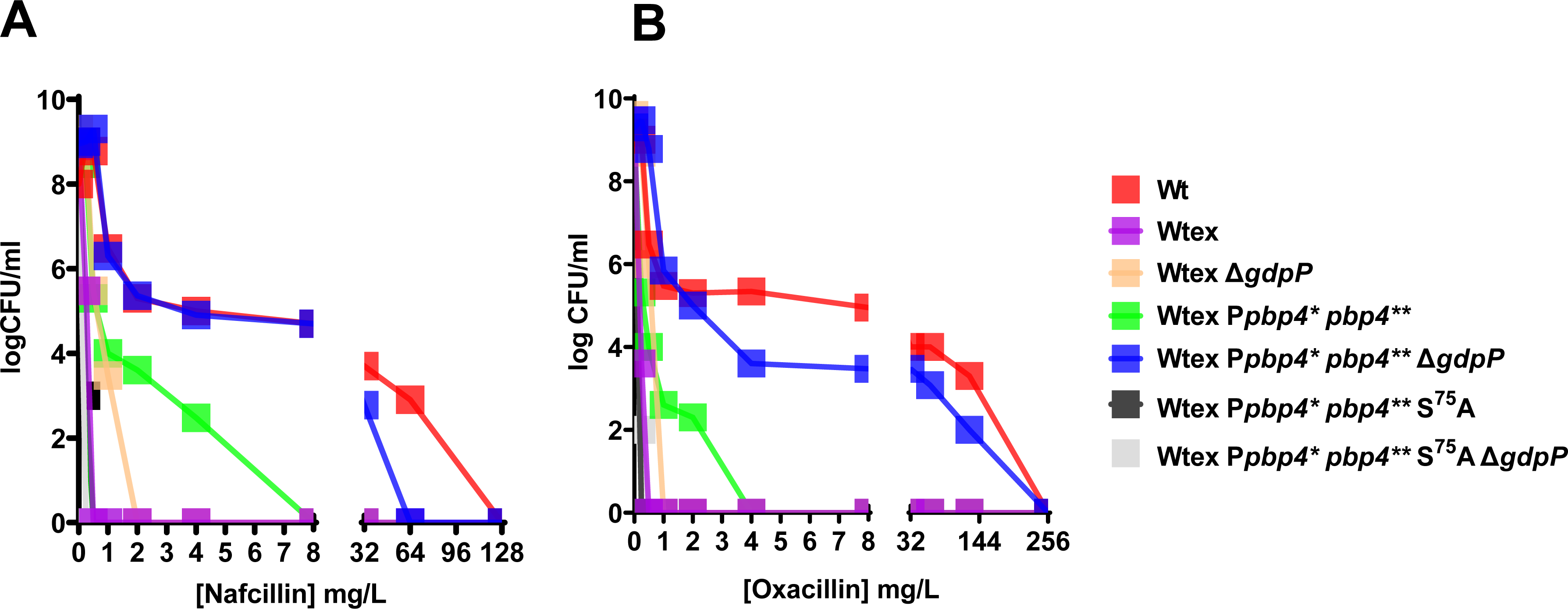
*pbp4* mutations and deletion of *gdpP* synergistically mediate high-level β-lactam resistance in *S. aureus*. Population analysis with **(A)** Nafcillin and **(B)** Oxacillin. The triple mutant, Wtex P*pbp4*\* *pbp4*\*\* Δ*gdpP* (blue squares) had increased survival compared to Wtex (pink squares), Wtex Δ*gdpP* (beige squares) and Wtex P*pbp4*\* *pbp4\*\** (green squares). Resistance displayed by the triple mutant was comparable to that displayed by Wt (red squares). Functional mutation of *pbp4* due to introduction of S^75^A (Wtex P*pbp4*\* *pbp4\*\** S^75^A, black squares and Wtex P*pbp4*\* *pbp4\*\** S^75^A Δ*gdpP*, grey squares) resulted in NGB susceptibility.

**Figure 4.**
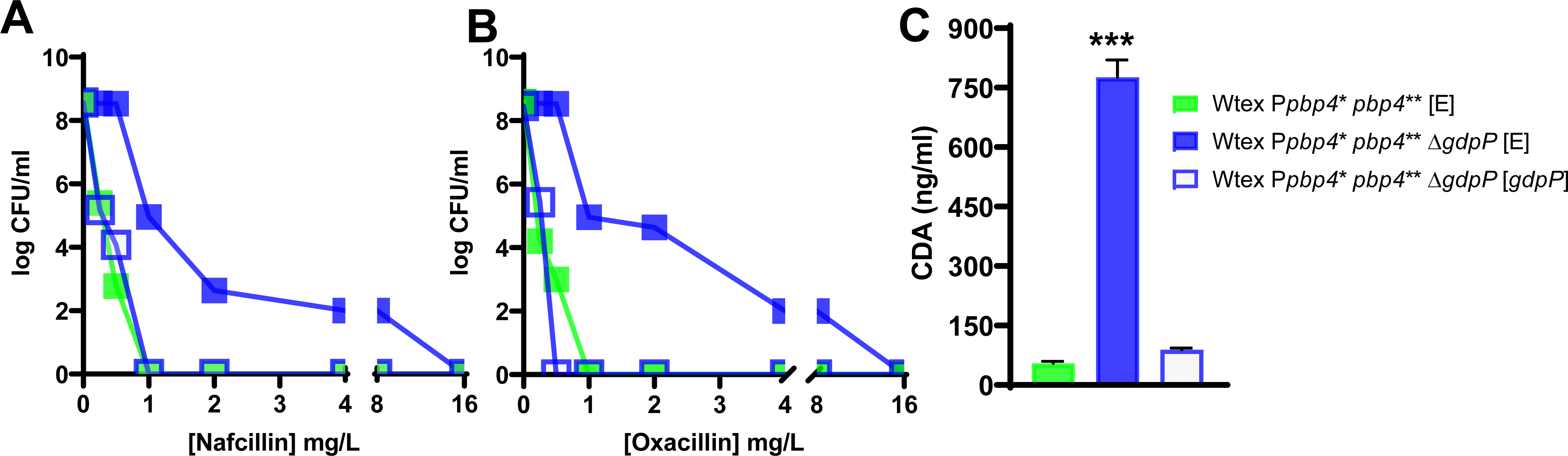
Complementation of *gdpP* restored β-lactam susceptibility of the NGB resistant triple mutant. Population analysis of complemented strains carried out with **(A)** Nafcillin and **(B)** Oxacillin. Complementation of the triple mutant with a functional GdpP (Wtex P*pbp4*\* *pbp4*\*\* Δ*gdpP* [*gdpP*], empty blue squares) resulted is a loss of high-level resistance, in a profile similar to Wtex P*pbp4*\* *pbp4*\*\* [E] (filled green squares). Complementation of the triple mutant with an empty vector (Wtex P*pbp4*\* *pbp4*\*\* Δ*gdpP* [E], filled blue squares) preserved the resistant phenotype associated with the mutations. **(C)** Measurement of intracellular levels of CDA for complemented strains. Complementation of the triple mutant with a functional GdpP (Wtex P*pbp4*\* *pbp4*\*\* Δ*gdpP* [*gdpP*], empty blue squares) led to decreased levels of CDA within cells that was similar to Wtex P*pbp4*\* *pbp4*\*\* [E] (filled green squares), whereas complementation of the triple mutant with an empty vector (Wtex P*pbp4*\* *pbp4*\*\* Δ*gdpP* [E], filled blue squares) maintained high levels of CDA.

**Table 1.**
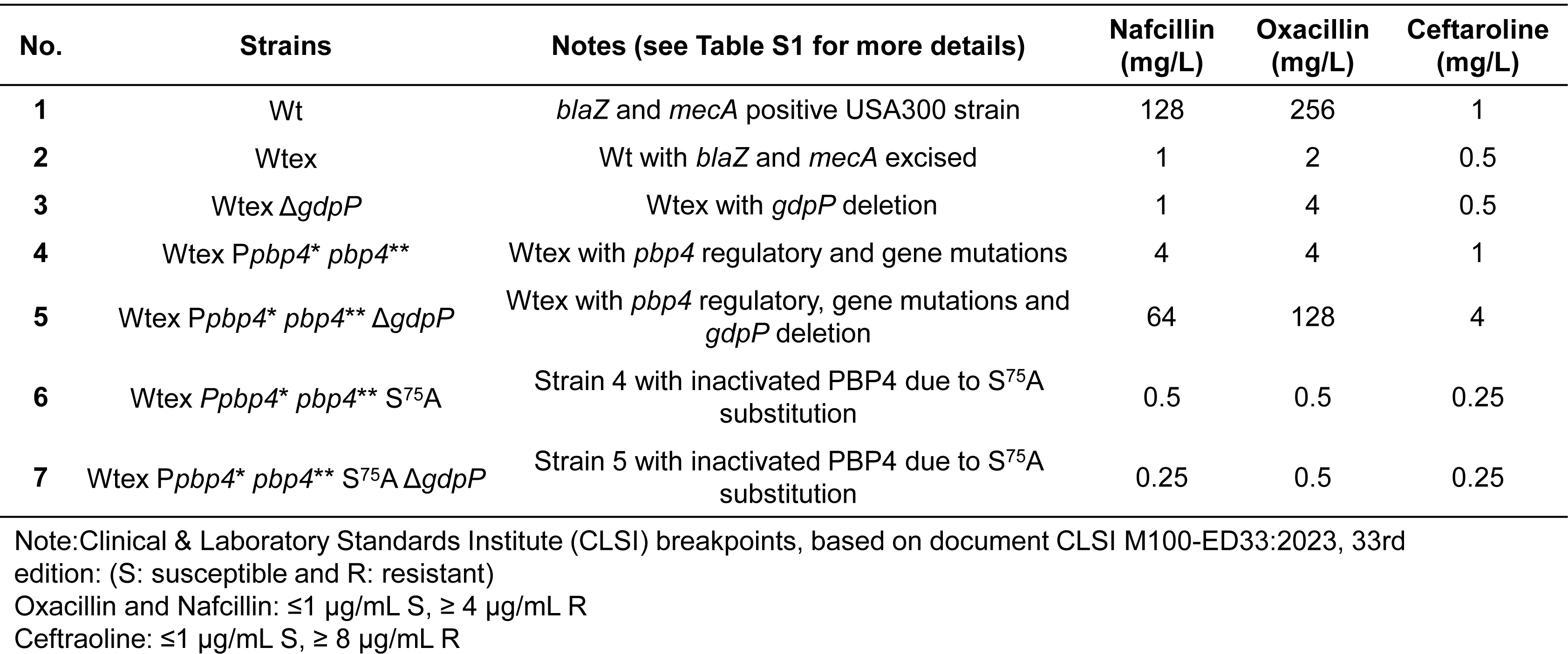
MIC assay of isogenic strains used in this study.

**Table 2.** MIC assay of complemented strains.

| Strains | Notes (see Table S1 for more details) | Nafcillin<br>(mg/L) | Oxacillin<br>(mg/L) | Ceftaroline<br>(mg/L) |
| --- | --- | --- | --- | --- |
| Wtex *Ppbp4\** *pbp4\*\** [E] | Strain 4 of Table 1 complemented with empty vector | 1 | 1 | 1 |
| Wtex *Ppbp4\** *pbp4\*\** Δ*gdpP* [E] | Strain 5 of Table 1 complemented with empty vector | 32 | 64 | 4 |
| Wtex *Ppbp4\** *pbp4\*\** Δ*gdpP* [*gdpP*] | Strain 5 of Table 1 complemented with constitutively expressing *gdpP* | 2 | 2 | 1 |

### Alterations in PBP4 and GdpP functions lead to synergistic increase in resistance, but maintain independent phenotypes

Since PBP4 and GdpP alterations resulted in a synergistic NGB resistance, we explored the mechanisms that could result in this synergy, and assessed whether either of the proteins imparted an influence over the functioning of the other. We first assessed whether the deletion of GdpP affected expression of PBP4, which in turn led to alterations in cell wall crosslinking and producing NGB resistance. Bocillin assay demonstrated that the expression of PBP4, along with the other PBPs, remained unaffected by *gdpP* deletion, as there was no difference seen in PBP4 levels among Wtex and Wtex Δ*gdpP*, or, Wtex P*pbp4*\* *pbp4*\*\* and Wtex P*pbp4*\* *pbp4*\*\* Δ*gdpP* strain pairs **(Fig 2D)**. Since the deletion of *gdpP* did not affect expression levels of PBPs, we next characterized the cell wall profiles of the strains to determine whether the mutations brought about any changes to the cell wall composition **(Fig 5)**. In alignment with the observation that *gdpP* deletion had no effect on PBP4 expression as seen in the Bocillin assay, the deletion of *gdpP* also did not have an effect on the cell wall composition (Fig 5A). There were no significant differences seen between the muropeptide profiles for Wt and Wtex, but the presence of *pbp4*- associated mutations, as seen previously (24), led to a significant increase in cell wall cross- linking as indicated by the increase in the 18+ muropeptide peaks, or the “hump” **(Fig 5B).** *pbp4*- associated increase in cell wall cross-linking remained unaltered in the presence of Δ*gdpP* in the triple mutant **(Fig 5C).** Taken together, these findings suggested that altered GdpP activity did not have any direct effect on the expression of PBPs, or the peptidoglycan composition.

**Figure 5:**
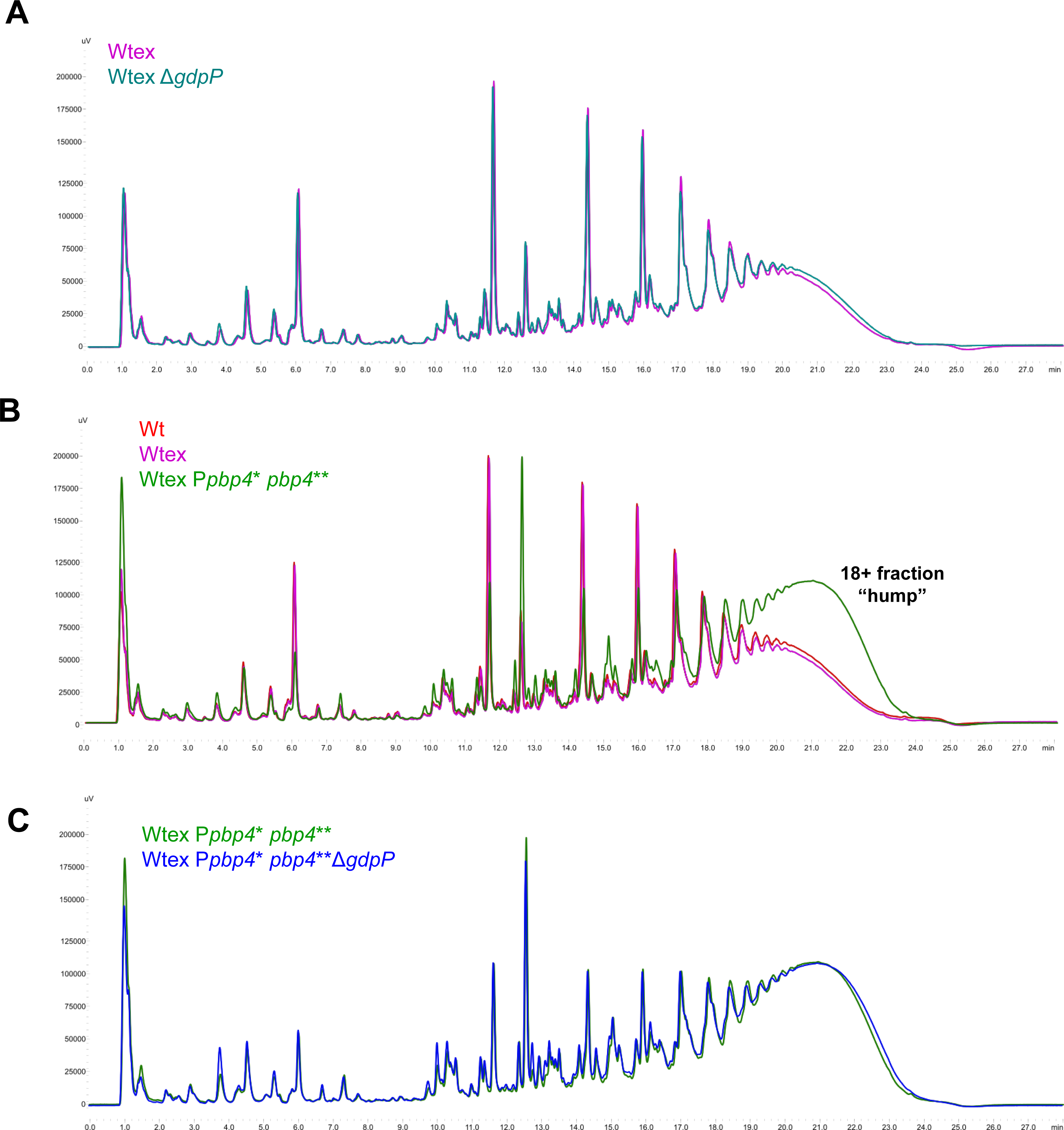
Alteration of GdpP does not affect the composition of the cell wall. Muropeptide purification and analysis profiles of **(A)** Wt, Wtex and Wtex P*pbp4*\* *pbp4*\*\* **(B)** Wtex and Wtex Δ*gdpP* **(C)** Wtex P*pbp4*\* *pbp4*\*\* and Wtex P*pbp4*\* *pbp4*\*\* Δ*gdpP. pbp4*-associated mutations led to increased cell wall cross-linking (increased levels of peak 12 and peak 18+ i.e. the “hump”) compared to Wtex. This increase was independent of Δ*gdpP*. Similarly, deletion of *gdpP* did not contribute to any significant alterations in the cell wall, irrespective of the presence of *pbp4*-associated mutations.

We next assessed if the *pbp4*-associated mutations affected GdpP-induced tolerance phenotype. A hallmark of *gdpP* deletion and the subsequent elevation in intracellular levels of CDA is the ability of cells to exert tolerance towards β-lactams (21). To determine this, a Tolerance-Disc (TD) test was performed **(Fig 6, Fig S5)** to measure the tolerance of each mutant to Nafcillin **(Fig 6A, 6B)** or Oxacillin **(Fig 6C, 6D)**. Relatively similar amounts of colony forming units (CFUs) were detected in both strains containing *gdpP* deletion (P value for Wtex Δ*gdpP* and Wtex P*pbp4*\* *pbp4*\*\* Δ*gdpP* = 0.7779 for Nafcillin and 0.2767 for Oxacillin; not significant) suggesting that the presence of *pbp4*-associated mutations did not affect drug tolerance. Strains with an unaltered GdpP, namely Wtex and Wtex P*pbp4*\* *pbp4*\*\* did not generate any tolerant colonies, suggesting that tolerance was attributed only to the deletion of *gdpP* **(Fig 6A, 6C)**. These findings suggested that *pbp4* and *gdpP* associated mutations independently led to β-lactam resistance and tolerance, respectively, and alterations in their functions did not impart any influence on the other.

**Figure 6.**
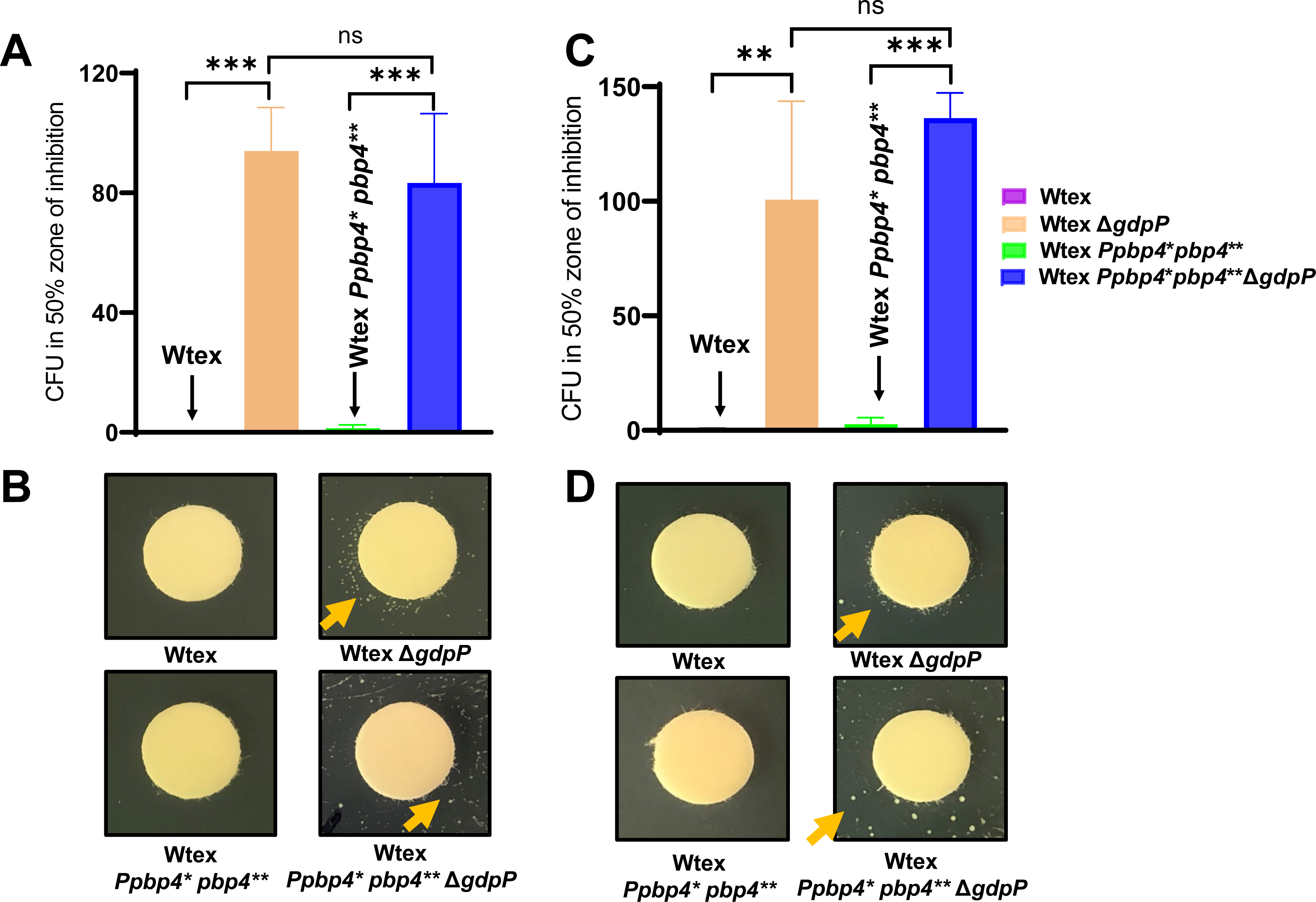
Deletion of *gdpP* led to NGB tolerance in *S. aureus.* TD-test analysis to estimate tolerance levels *in S. aureus* strains. **(A-B)** TD-test with Nafcillin. **(C-D)** TD-test with Oxacillin. CFUs in the 50% inhibition zone of the strains containing *gdpP* deletion, namely Wtex Δ*gdpP* and Wtex P*pbp4*\* *pbp4*\*\* Δ*gdpP* are higher compared to their isogenic strains with a functional GdpP, namely Wtex and Wtex P*pbp4*\* *pbp4*\*\*. All data are from three independent experiments and presented as mean ± SD with One-Way ANOVA (***: P-value ≤ 0.001, **: P-value ≤ 0.01).

### Synergistic action of PBP4 and GdpP alterations resulted in NGB therapy failure

To determine if synergistic action of altered functioning of PBP4 and GdpP, which brought about high-level resistance to NGBs *in vitro* had an effect on bacterial survival during *in vivo* infection, we performed infection assays with *C. elegans* **(Fig 7A to 7D)**. Infection was performed with Wt, Wtex or Wtex P*pbp4*\* *pbp4*\*\* Δ*gdpP* in presence of increasing concentrations of Nafcillin. While worms infected with Wtex had maximum survival, infection with Wtex P*pbp4*\* *pbp4*\*\* Δ*gdpP* led to significant worm killing even in presence of 4 mg/L Nafcillin (80% worm survival) **(Fig 7A)** or 8 mg/L Nafcillin (85% worm survival), suggesting that the triple mutant maintained its resistant phenotype during *in vivo* infection **(Fig 7B)**. The worm survival was seen to be over 90% for all three strains only at the high concentration of Nafcillin at 16 mg/L. **(Fig 7C)**. The killing pattern for the triple mutant was very similar to that displayed by Wt, which also resulted in killing of worms despite the presence of Nafcillin **(Fig 7A, 7B)**, and required high concentrations of Nafcillin in order to result in >90% worm survival **(Fig 7C)** thus reiterating the MRSA-like resistance phenotype of the triple mutant. In the absence of antibiotics, Wt and Wtex strains led to maximum killing of the worms, resulting in 35% survival at the end of the 72h assay. However, the triple mutant had significantly attenuated survival, where worm survival was over 70%, suggesting an attenuation in virulence **(Fig 7D)**. OP50, the *E. coli* control, maintained a 100% survival rate throughout the assay, indicating that the killing observed was attributed to *S. aureus* strains.

**Figure 7.**
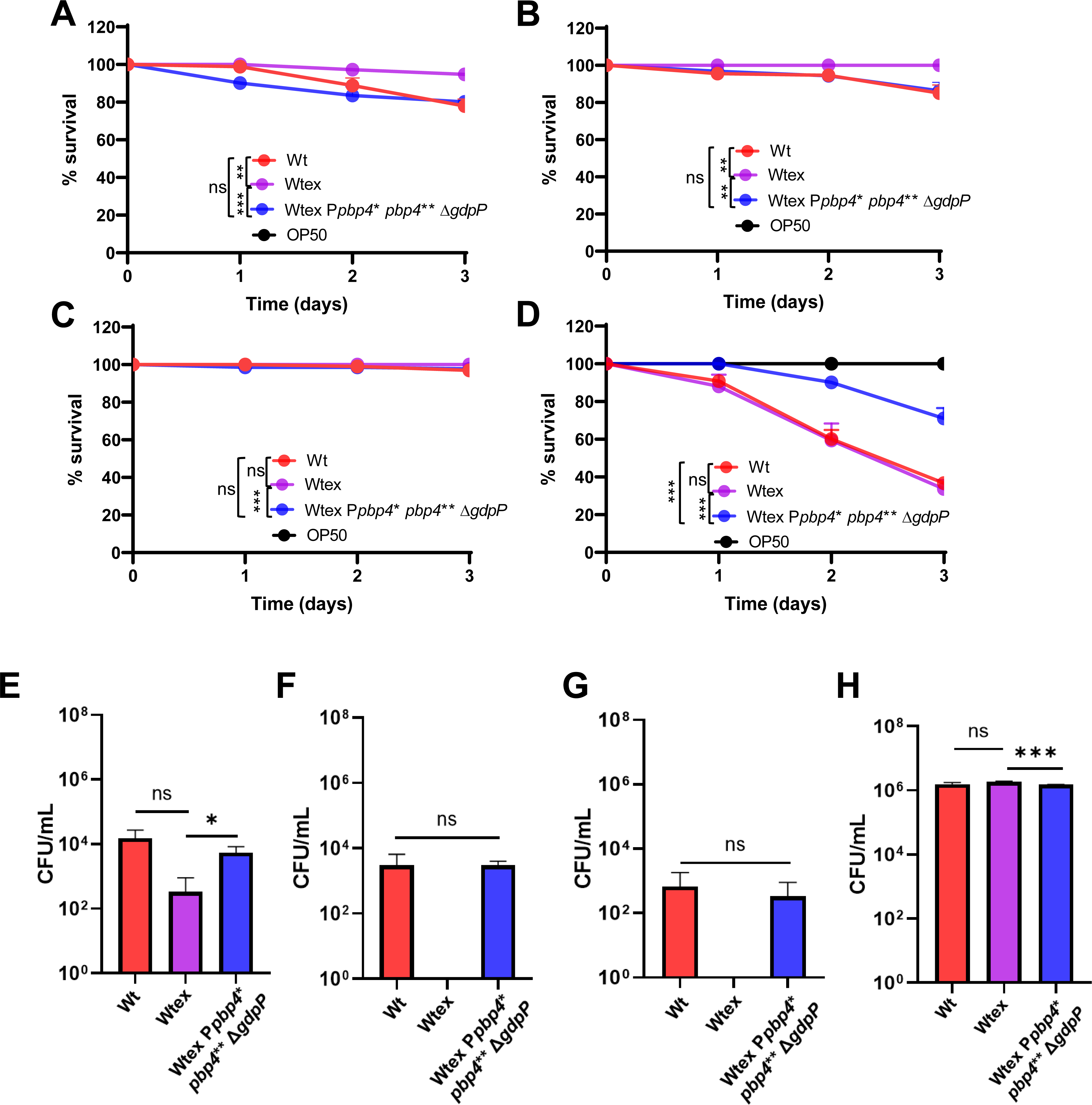
Synergistic action of PBP4 and GdpP alterations resulted in NGB therapy failure. Percent survival of *C. elegans* when infected with bacteria over a period of 72h in presence of **(A)** 4 mg/L **(B)** 8 mg/L or **(C)** 16 mg/L Nafcillin. Worms infected with Wtex had 100% survival, whereas infection with Wtex P*pbp4*\* *pbp4*\*\* Δ*gdpP* had decreased survival, similar to that seen by Wt. **(D)** The triple mutant, Wtex P*pbp4*\* *pbp4*\*\* Δ*gdpP* had attenuated virulence compared to Wt and Wtex the the absence of Nafcillin. Worms infected with OP50, the *E. coli* control, showed 100% worm survival. Enumeration of bacterial load within the gut of *C. elegans* in presence of **(E)** 4 mg/L **(F)** 8 mg/L or **(G)** 16 mg/L Nafcillin. Compared to Wtex, which was was not detected, Wt and Wtex P*pbp4*\* *pbp4*\*\* Δ*gdpP* had significant colonization within the gut. **(H)** Gut bacterial enumeration when infection was carried out without Nafcillin indicated that Wt, Wtex and Wtex P*pbp4*\* *pbp4*\*\* Δ*gdpP* had similar colonization abilities. (***: P-value ≤ 0.001, and **: P-value ≤ 0.01, P-value ≤ 0.05, ns = P-value > 0.05)

The bacterial load within the *C. elegans* gut was determined at the end of the assay by lysing the worms and enumerating CFUs **(Fig 7E-7H)**. When infection was performed in presence of Nafcillin, CFU counts were the highest for Wt, followed by the triple mutant in a concentration dependent manner, indicating that Wt and Wtex P*pbp4*\* *pbp4*\*\* Δ*gdpP* were successfully able to colonize *C. elegans* despite the presence of Nafcillin **(Fig 7E-7G)**. No CFUs were detected for Wtex, which was in alignment with 100% worm survival. In case of infection without Nafcillin, there was no significant difference in CFUs among the strains, suggesting similar levels of bacterial colonization **(Fig 7H)**. Taken together, these findings demonstrated that the triple mutant, similar to Wt, could successfully colonize and kill *C. elegans* in presence of Nafcillin, suggesting that strains with alterations in PBP4 and GdpP have increased fitness during *in vivo* infection, and can lead to therapy failure when treated with NGBs, in a manner similar to that seen by MRSA strains.

## Discussion

*S. aureus* is a frequent colonizer in humans, and a major cause of infections associated with skin and soft tissue, lower respiratory tract, blood or bones and joints (1, 37). Treatment of *S. aureus* can be challenging not only due to the nature of the infection that it causes, which ranges from bacteremia to biofilm formation, but also due to the ability of the pathogen to evade antibiotic treatment (2). Resistance to safe and effective antibiotics such as Next Generation β-lactams (NGBs) compels clinicians often to opt for alternatives that are known for having lower efficacy and adverse side effects (38). For the past several decades, *mecA or mecC* have been the hallmark of high-level NGB resistance in *S. aureus*, thus restricting the use of NGBs for treatment (8). However, NGBs (such as Nafcillin and Cefazolin) remain as the go-to drug for treatment of infections caused by *S. aureus* strains that lack *mecA* (termed as MSSA), which are currently being detected in increasing numbers worldwide (5). Despite lacking *mecA*, certain strains of MSSA can display NGB resistance and are termed as MRLMs (<u>M</u>ethicillin <u>R</u>esistance <u>L</u>acking *<u>m</u>ec*) making Nafcillin or Cefazolin’s use ineffective. Following early identification in the 1980’s, MRLMs have also been detected in recent years (13-15) **(Fig 1)**. MRLMs can be misdiagnosed as MSSA due to the lack of *mec* genes, which may cause improper therapeutic intervention, longer hospitalization and increased healthcare related cost. The underlying basis of MRLMs’ resistance to NGB remained unknown. In this study, we demonstrated that mutations that alter PBP4 expression and function along with mutations in *gdpP* that increase CDA production, can synergistically mediate NGB resistance **(Fig 3**, **Fig 8, Fig S4)**. Abolition of function of either PBP4 (through S75A substitution at its active site) or reduction of CDA concentrations (through complementation with *gdpP*) produced total NGB susceptibility in the resistant strain. These results suggested that both PBP4 and CDA played an equally important role that caused NGB resistance in the resistant strain. To our surprise, NGB resistance in the triple mutant, P*pbp4*\* *pbp4*\*\* Δ*gdpP*, was not only significantly higher than its parental strain Wtex, but was comparable to Wt, a MRSA strain. Notably, the triple mutant out-completed Wt when challenged with Ceftaroline (a highly advanced NGB that is currently used to treat complicated MRSA infections) when assessed through MIC assay **(Table 1**, **Table 2)**. To assess these results more objectively, we also performed growth curve analysis with Ceftaroline, which produced identical results to that of MICs **(Fig 8)**. Thus, the synergistic mechanism of NGB resistance presented in this study not only identifies as at least one of the basis through which MRMLs could arise but also suggests that this mode of resistance is highly effective in producing NGB resistance that is typically observed in MRSAs. Furthermore, the *in vivo* experiments demonstrated that in the presence of Nafcillin, the NGB-resistant triple mutant, was able to kill *C. elegans* at a significantly higher level compared to its NGB susceptible parental strain **(Fig 7)**, indicating the possibility of therapy failure, or an increase in therapeutic complications. Infection of *C. elegans* in the absence of drug depicted a significant decrease in virulence of the triple mutant. This phenotype was consistent with previous studies demonstrating decreased *C. elegans* infection by strains possessing *pbp4*- associated mutations (28). Further, accumulation of intracellular CDA has also been previously associated with decreased virulence (39, 40). In line with our previously published results, strains with *gdpP* deletion, including the triple mutant, displayed decreased hemolysis when plated onto blood-agar plates suggesting decreased virulence **(Fig S6)**, and reiterating that alterations associated with *pbp4* and *gdpP* alterations maintained their well-characterized, independent phenotypes in the triple mutant.

**Figure 8.**
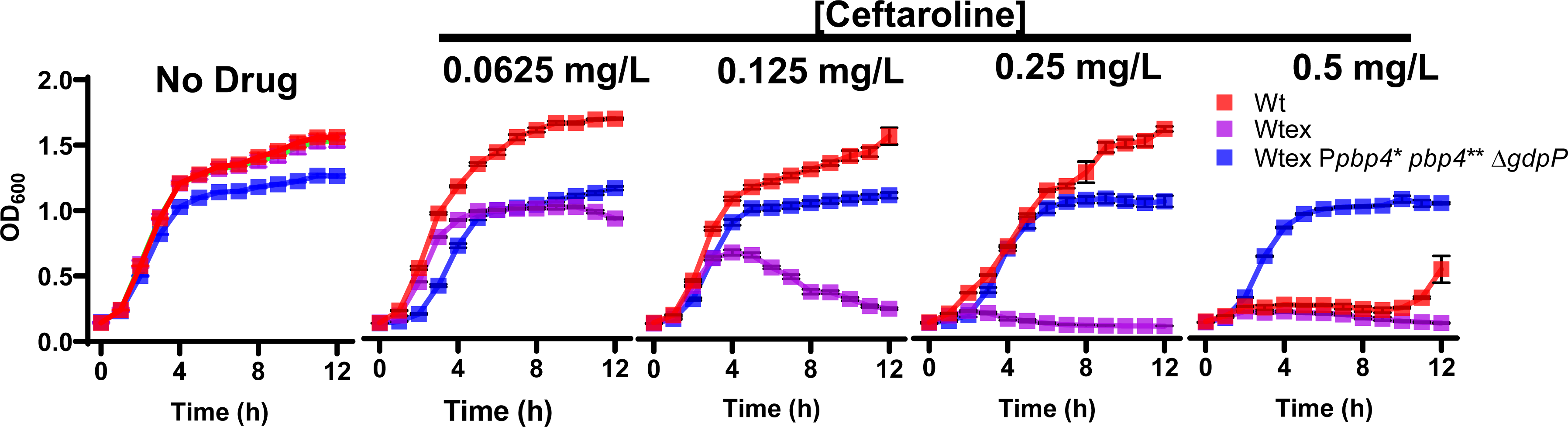
Growth assay in presence of Ceftaroline. Treatment of Wt, Wtex and Wtex P*pbp4*\* *pbp4*\*\* Δ*gdpP* in increasing concentrations of Ceftraroline (0-0.5 mg/L) demonstrated that Wt and Wtex were susceptible to Ceftaroline. However, and Wtex P*pbp4*\* *pbp4*\*\* Δ*gdpP* was not susceptible to the drug and survived even in presence of the highest concentration of the drug.

The results of our study also allude to a potential interaction between pathways involving PBP4 and GdpP that results in NGB resistance. PBP4 over-expression results in increased peptidoglycan cross-linking (24); we thus speculated if increased CDA levels led to an increase in the expression of PBPs that potentiated cell-wall crosslinking. However, Bocillin assay and peptidoglycan analysis indicated that increased CDA did not have any effect on the expression of PBPs, or alter the cell wall profile, suggesting that there was likely no canonical change in the cell wall cross-linking in the triple mutant that facilitated the resistant phenotype **(Fig 2D**, **Fig 5)**. Similarly, our results also indicated that PBP4 overexpression did not influence tolerance to NGBs, a phenotype typically associated with increased CDA concentrations. Thus, the mechanism of synergism that brings about NGB resistance remains unclear at this point. We hypothesize that increased CDA concentrations influence PBP4 localization, stability or turnover either in a direct or indirect manner to mediate high-level NGB resistance synergistically. In bacteria including *S. aureus*, *Listeria monocytogenes* and *Bacillus subtilis*, CDA has been associated with maintenance and stability of cell wall and β-lactam resistance (41-43). CDA also regulates the uptake of potassium ions by interacting with transporters such as KtrAB/AD and KdpFABC (44). In *Listeria monocytogenes*, CDA accumulation led to impaired activity of the D- alanine ligase, *ddl* due to decreased uptake of potassium ions, resulting in decreased level UDP- N-acetylmuramic acid and impaired peptidoglycan synthesis (45). However, such an alteration would have resulted in an altered muropeptide profile for the Δ*gdpP* strains (Fig 5A). Conversely, in *Lactococcus lactis*, accumulation of CDA led to increased intracellular levels of UDP-N- acetylglucosamine, an integral component of the peptidoglycan (46, 47). Taken together, multiple studies have drawn associations between CDA and peptidoglycan synthesis, but it is evident that these associations are via intricate pathways, and may vary between bacterial species. Thus, further studies are necessary to elucidate the role of CDA levels in peptidoglycan synthesis.

In summary, this study highlights that PBP4 and GdpP are together, important players of NGB resistance and could have important roles in the development of MRLM strains, and should be further studied in order to uncover the mechanistic pathway leading to the synergistic, high-level NGB resistance.

## Acknowledgement

Research reported in this publication was supported by the National Institute of Allergy And Infectious Diseases of the National Institutes of Health under Award Number R01AI165510 & 2R01AI100291 to SSC. The content is solely the responsibility of the authors and does not necessarily represent the official views of the National Institutes of Health. SSC would also like to acknowledge the University of Maryland, Baltimore, and the University of Maryland Center for Environmental Science for providing seed funds. NS received Student Enhancement Funds from the Institute of Marine and Environmental Technology (IMET) that partially covered travel expenses to Portugal for performing the cell wall analysis. LBM was funded by fellowship UI/BD/153384/2022 from Fundação para a Ciência e a Tecnologia. The authors would like to thank Dr. Nicholas Carbonetti for his feedback on this manuscript.

